# Environment-dependent epistasis increases phenotypic diversity in gene regulatory networks

**DOI:** 10.1101/2022.09.18.508240

**Authors:** Florian Baier, Florence Gauye, Ruben Perez-Carrasco, Joshua L. Payne, Yolanda Schaerli

**Affiliations:** Department of Fundamental Microbiology, University of Lausanne, Biophore Building, 1015 Lausanne, Switzerland; Department of Life Sciences, Imperial College London, London SW7 2AZ, UK; Institute of Integrative Biology, ETH Zurich, 8092 Zurich, Switzerland; Swiss Institute of Bioinformatics, 1015 Lausanne, Switzerland

## Abstract

Mutations to gene regulatory networks can be maladaptive or a source of evolutionary novelty. Epistasis confounds our understanding of how mutations impact the expression patterns of gene regulatory networks, because such nonlinearities make it difficult to predict the combined phenotypic effects of mutations based on knowledge of the mutations’ individual effects. This challenge is exacerbated by the dependence of epistasis on the environment, which is particularly germane to gene regulatory networks that interpret signals in space or time. To help fill this knowledge gap, we used the toolkit of synthetic biology to systematically assay the effects of pairwise and triplet combinations of mutant genotypes on the expression pattern of a gene regulatory network expressed in *Escherichia coli* that interprets an inducer gradient across a spatial domain. We uncovered a preponderance of epistasis in both pairwise and triplet combinations that can switch in magnitude and sign across the inducer gradient to produce a greater diversity of expression pattern phenotypes than would be possible in the absence of such environment-dependent epistasis. We discuss our findings in the context of the evolution of hybrid incompatibilities and evolutionary novelties, arguing that environment-dependent epistasis is likely an important cause of both phenomena in gene regulatory networks.

## Introduction

The regulation of gene expression is essential for the spatiotemporal control of diverse biological functions. Gene regulation is mainly mediated by *trans*-regulatory proteins and protein complexes such as transcription factors and RNA polymerase, which target specific DNA sequences in *cis*-regulatory regions such as promoters and enhancers to modulate gene expression levels (Signor and Nuzhdin, 2018). Transcription factors often regulate their own expression levels, as well as the expression levels of other transcription factors, giving rise to gene regulatory networks (Alon, 2007). Gene regulatory networks drive fundamental physiological and developmental processes, such as the interpretation of morphogen gradients for spatial patterning during embryogenesis (Ben-Tabou de-Leon and Davidson, 2007).

Given their central role in essential biological functions, it is crucial that gene regulatory networks are robust to genetic perturbation. Indeed, mutations in *cis*-regulatory regions that induce quantitative (e.g., DNA mutations that alter the affinity of a transcription factor binding site) or qualitative (e.g., DNA mutations that create or destroy a transcription factor binding site) changes to a gene regulatory network often do not change the network’s spatiotemporal expression pattern phenotype (Dalal and Johnson, 2017; Payne and Wagner, 2015). This robustness causes gene regulatory networks to be highly evolvable, because it facilitates the neutral accumulation of mutations (Ciliberti et al., 2007; Cotterell and Sharpe, 2010; Payne and Wagner, 2019; van Nimwegen et al., 1999). This creates genetic diversity and promotes “system drift” (True and Haag, 2001), in which a population undergoes a series of quantitative and qualitative changes to a gene regulatory network that are phenotypically neutral (Crombach et al., 2016; Dalal and Johnson, 2017). Subsequent mutations to, or recombination events among, such diverse network configurations can then generate phenotypic variation (Ciliberti et al., 2007; Johnson, 2017; Martin and Wagner, 2009).

Mutations to gene regulatory networks are commonly implicated in evolutionary adaptations and innovations (Johnson, 2017; Prud’homme et al., 2007). There has been an intense research effort to understand the molecular details of these mutations and the mechanistic basis of how they alter gene expression pattern phenotypes (from here on referred to as pattern phenotypes) (Nghe et al., 2020). A common observation is that the combination of two (or more) mutations can result in phenotypic effects that would not be expected based on an additive assumption of each single mutant’s phenotypic contribution (Lagator et al., 2017b; Li et al., 2019; New and Lehner, 2019). Such context-dependence of mutational effects is called epistasis and is referred to as negative (positive) when the combined effects of mutations are less than (more than) expected based on their individual effects. Moreover, epistasis can itself be dependent on environmental conditions, such as the concentration of an expression inducer or an enzymatic co-factor (de Vos, Poelwijk et al. 2013, Flynn, Cooper et al. 2013, de Vos, Dawid et al. 2015, Lagator, Sarikas et al. 2017, Li and Zhang 2018, Li, Lalic et al. 2019, Anderson, Baier et al. 2021). For example, in the lambda phage promoter, a canonical gene regulatory system, Lagator and colleagues (2017a) found that 67% (14%) of 141 double mutants exhibited negative (positive) epistasis when the transcription factor that competes for binding with RNA polymerase was not expressed, and that 58% of the double mutants switched from negative to positive epistasis (or vice versa) when the transcription factor was expressed. Epistatic interactions and their dependence on the environment are not limited to pairs of mutations, but can also occur amongst three or more mutations (Domingo et al., 2018; New and Lehner, 2019), a phenomenon known as higher-order epistasis (Weinreich et al., 2013).

Epistasis in GRNs has many causes including specific interactions across intermolecular interfaces, such as protein-protein (Diss and Lehner, 2018; Podgornaia and Laub, 2015) and transcription factor-DNA interactions (Anderson et al., 2015). This is the particular case of typical non-linearities inherent to competitive and cooperative binding of several transcription factors to the same target. In addition, the presence of feedback and feedforward loops in the network introduce additional epistatic effects that depend on the topology of the GRN (Domingo et al., 2019; Nghe et al., 2020).

Despite substantial progress in the characterization of epistasis in macromolecules (Julien et al., 2016; Li et al., 2016; Olson et al., 2014; Puchta et al., 2016) and gene regulatory networks (Lagator et al., 2017a; Lagator et al., 2017b; Schaerli et al., 2018), it remains poorly understood how environment-dependent epistasis influences the spatiotemporal pattern phenotypes of gene regulatory networks. This is an important knowledge gap, because epistasis can constrain or facilitate evolvability (Nghe et al., 2020; Payne and Wagner, 2019), and the interpretation of chemical gradients (i.e., the environment) by gene regulatory networks is fundamental to developmental patterning (Ashe and Briscoe, 2006; Rogers and Schier, 2011), as well as other essential biological processes like chemotaxis (Swaney et al., 2010). One of the reasons this knowledge gap persists is the difficulty of studying gene regulatory networks *in situ*, due to their embedding in large and complex cellular networks.

Synthetic biology offers a path forward (Baier and Schaerli, 2021; Crocker and Ilsley, 2017). By extracting a gene regulatory network from its native cellular environment, it becomes feasible to systematically study the effects of mutations, individually and in combination, on the pattern phenotype of a gene regulatory network. We recently built and studied a synthetic three-node gene regulatory network, expressed in *Escherichia coli (E. coli)*, that produces a stripe pattern phenotype (low-high-low) along an inducer gradient – analogous to a morphogen gradient interpreted during embryogenesis (Schaerli et al., 2014). The network topology was based on the incoherent feed-forward loop 2 motif (Alon, 2007), which drives numerous biological functions, such as blastoderm patterning in *Drosophila* (Jaeger et al., 2012). We previously used this synthetic gene regulatory network to study how mutation brings forth phenotypic variation in the network’s pattern phenotype (Schaerli et al., 2018).

Here, we systematically combined mutations in the cis-regulatory regions of each of the network’s three nodes, covering promoters and transcription factor binding sites, into pairwise and triplet combinations. We analyzed how pairwise and higher-order epistasis varies along the inducer gradient. In doing so, we provide the first study of how environment-dependent epistasis influences the pattern phenotype of a gene regulatory network. Our results reveal a context-dependent picture of epistasis in which genotype combinations exhibit diverse epistatic effects, including positive and negative epistasis, that can change drastically along the inducer gradient. In turn, this inducer dependence of epistasis leads to more diverse pattern phenotypes in our network than would be expected if the mutations did not interact epistatically or depend on the environment. In the context of evolution, such increased diversity could facilitate adaptive evolution, but may also underlie hybrid incompatibility and speciation.

## Results

### Experimental system and approach

We studied a synthetic gene regulatory network with three nodes, which we refer to as sensor, regulator and output (**Figure 1a)** (Schaerli et al., 2018; Schaerli et al., 2014). Given the correct promoter activity and repression between nodes, the network produces a stripe pattern phenotype (low-high-low gene expression of the output node) along an inducer gradient, using the following mechanism (**Figure 1b)**: The sensor is activated by the inducer and represses the regulator and the output. The regulator also represses the output, but its activity decreases with increasing inducer concentration, due to repression by the sensor. Consequently, the output is the least repressed at intermediate inducer levels, resulting in high expression and the formation of a stripe along the inducer gradient. Our definition of a stripe pattern phenotype requires the output expression to be the highest at intermediate inducer levels but leaves room for variation in terms of the stripe shape, intensity and overall expression (Schaerli et al., 2018).

**Figure 1.**
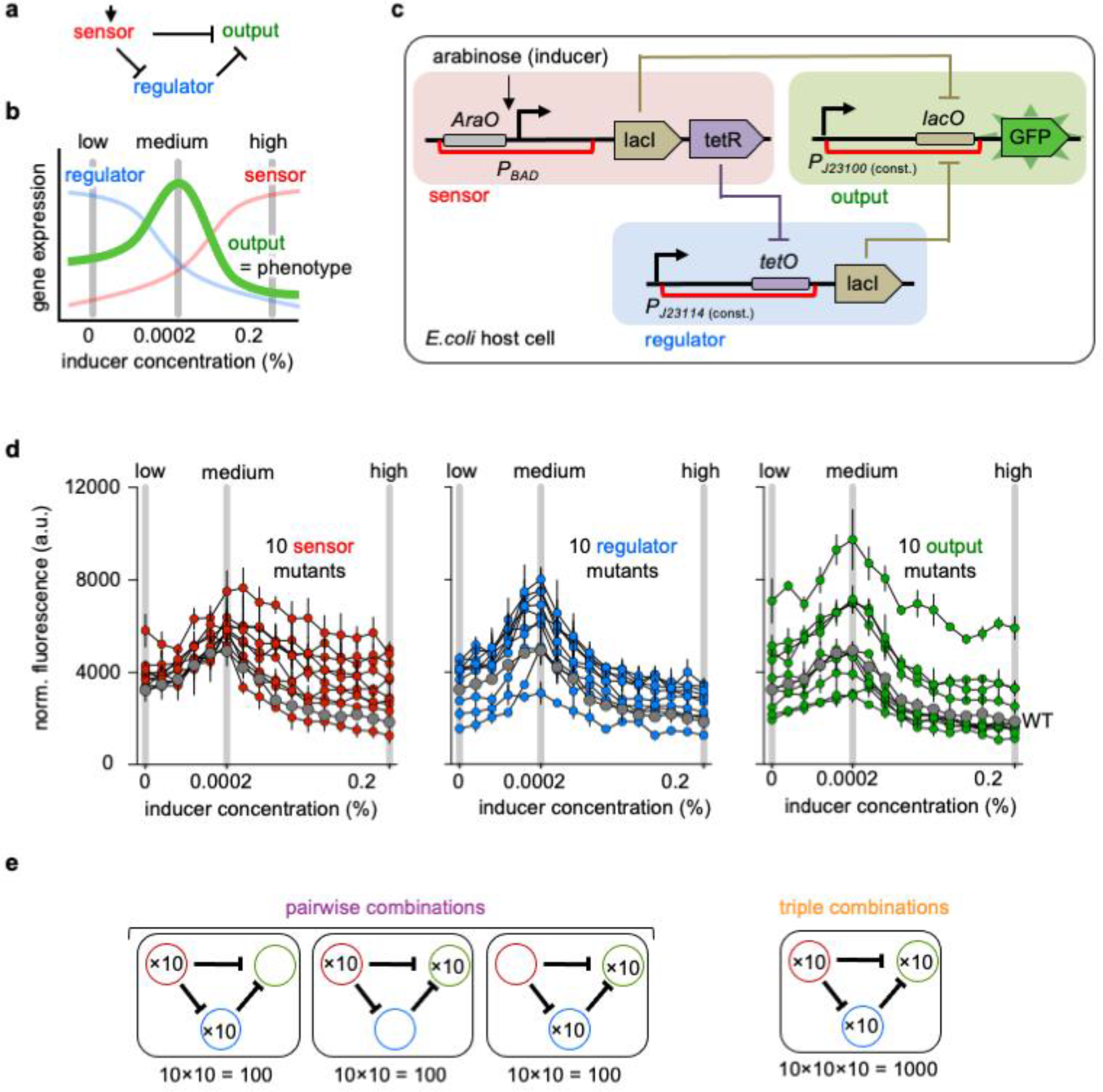
Experimental system. (**a**) Topology of the studied gene regulatory network. (**b**) Schematic of gene expression patterns of sensor, regulator and output nodes along an inducer gradient. The pattern phenotype of the “wild-type” (WT) network is a stripe of gene expression of the output node (green) along an inducer gradient. (**c**) Molecular implementation of the synthetic gene regulatory network in E. coli. The network is induced with arabinose through a pBAD promoter. Repressive regulatory interactions are implemented with LacI and TetR repressor proteins binding to their respective operators lacO and tetO, which lowers transcription from the promoter upstream. The detected network output is fluorescence of GFP. The cis-regulatory regions containing mutations are indicated with red brackets. (**d**) GFP expression patterns of the 30 genotypes carrying mutations in the sensor (left), regulator (middle) or output (right) node. GFP expression of the WT network is shown in grey. Each genotype was measured in triplicate at 16 inducer concentrations and the mean and standard deviation from three biological replicates are shown. **(e)** Schematic of how we generated all 300 pairwise and 1000 triplet combinations.

We encoded the three nodes of the synthetic gene regulatory network on separate plasmids (one per node), which we expressed in *E. coli*. The arabinose-responsive promoter pBAD receives the inducer signal (arabinose) (**Figure 1c**). The inhibitions are implemented by the transcriptional repressors TetR (tetracycline repressor) and LacI (lactose repressor) binding to their operator sites (TetO and LacO), which we placed downstream of the regulator and output promoters, respectively. The observable network output is expression of the superfolder green fluorescent protein (GFP) from the output node (Pedelacq et al., 2006).

We previously introduced random nucleotide changes in the *cis*-regulatory regions spanning the promoter and operator sequences separately in each of the network nodes (Schaerli et al., 2018). From this study, we selected 31 genotypes: the starting network (“wild-type”, WT) and ten mutant genotypes for each of the three network nodes. Each mutant genotype contained one to three nucleotide changes in its *cis*-regulatory regions in only one of the nodes, either sensor, regulator or output (**Extended Figure 1.1**). All of them displayed a stripe phenotype, although with quantitative variations in overall fluorescence level and shape of the stipe pattern (**Figure 1d**). We refer to the 30 mutant genotypes as sensor-1, regulator-1, output-1, sensor-2, etc. The variation in GFP expression along the gradient reflects the mechanistic role of the mutated node in the network. For example, sensor genotypes showed high variation at higher inducer concentrations, caused by a weaker promoter and/or lower sensitivity towards the inducer, which resulted in a lower repression of the output node at high inducer concentrations. In contrast, output genotypes showed variation along the gradient, which is caused by changes in their promoter and operator strengths.

To study two-way and three-way epistasis between genotypes, we systematically combined all 3×10 mutant genotypes to generate all possible 300 (3×10×10) pairwise and 1000 (10×10×10) triplet combinations by transforming the plasmid combinations into *E. coli* cells (**Figure 1e**). We cultured cells in 384-well plates and measured GFP expression of all 300 pairwise and 1000 triplet genotypes in triplicate at low (0%), medium (0.0002%) and high (0.2%) inducer concentrations using a fluorescence spectrophotometer (**Figure 1b**). Fluorescence measurements correlated well between replicates (**Extended Figure 1.2a**, R^2^ of 0.90 for all pairwise and 0.95 for all triplet combinations measurements). The standard deviation of measurement was similar between single, pairwise and triplet genotypes (**Extended Figure 1.2b)**. In addition to the measurements at three inducer concentrations, we also assayed all single mutant genotypes (i.e. the 30 genotypes selected from (Schaerli et al., 2018)) and 40 pairwise and triplet genotypes at 16 inducer concentrations. These measurements over 16 inducer concentrations confirmed that the measurements at three inducer concentrations capture the pattern phenotypes well (**Extended Figure 1.2c and d**).

### Most genotype combinations exhibit epistasis

We used a multiplicative model to determine whether pairwise or higher-order interactions exhibited epistasis. In this model, epistasis is defined as the deviation of the observed gene expression of the pairwise or triplet combinations from the product of the single mutant expression levels relative to the WT genotype. The multiplicative model is commonly used to detect epistasis in gene regulatory systems (Lagator et al., 2017a; Lagator et al., 2017b; Li et al., 2019; New and Lehner, 2019), as well as in other systems (Gao et al., 2010; Poelwijk et al., 2019) (**Figure 2a**). The multiplicative model is also referred as log-additive, since it becomes additive when the concentrations are transformed to logarithmic scale.

**Figure 2.**
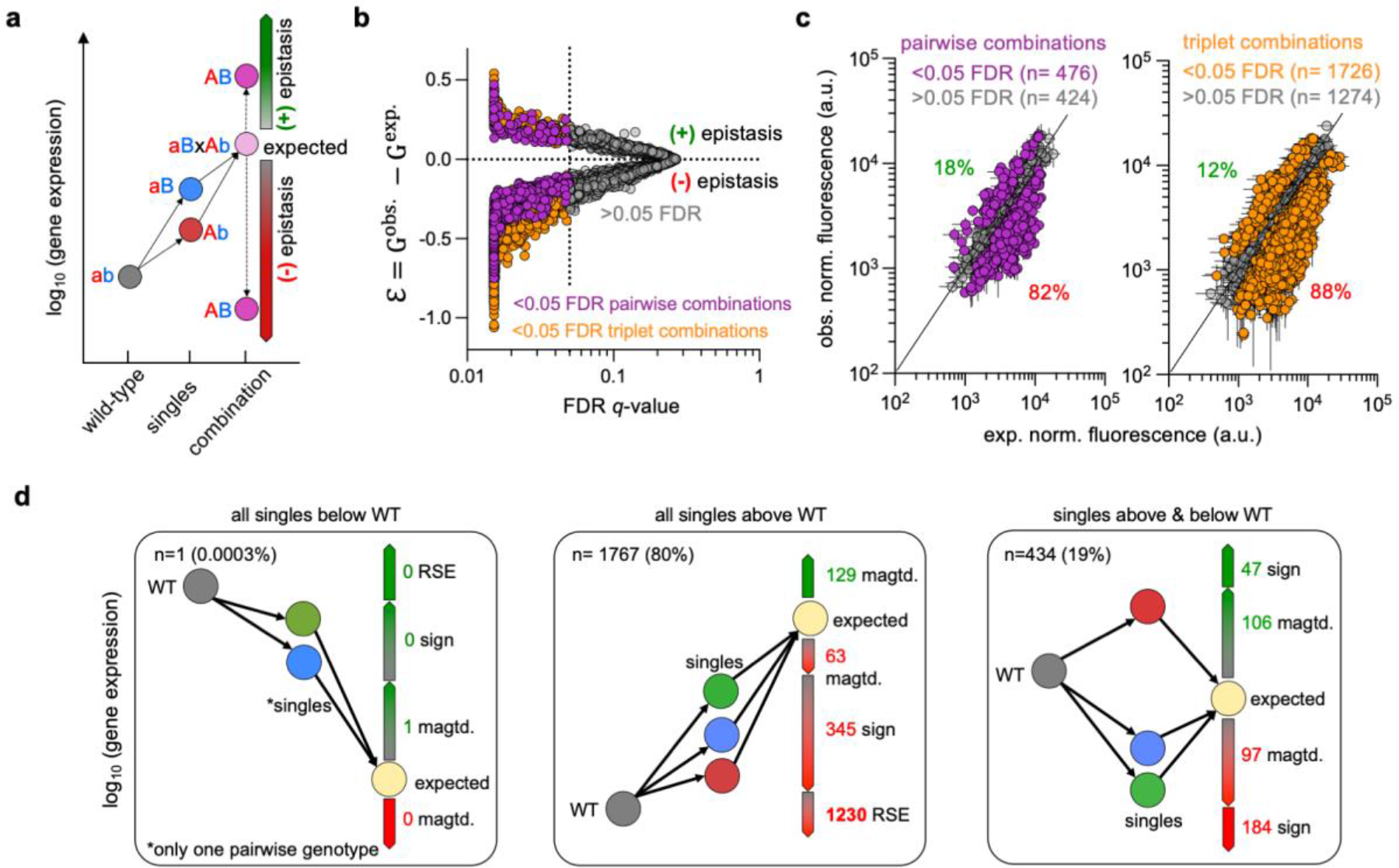
Definition and prevalence of epistasis. (**a**) Illustration of the multiplicative (log-additive) model for pairwise mutational combinations in a gene expression system. In this example, the gene expression of each single mutant genotype is higher than the WT (ab). If the observed gene expression of the pairwise genotype combination is significantly higher than what would be expected based on the single mutant genotypes, we define it as positive epistasis. If the observed gene expression is lower, we define it as negative epistasis. (**b**) Epistasis values and corresponding q-values, with significant values (FDR <0.05) in colour and non-significant values (FDR >0.05) in grey. Data from measurements at low (0%), medium (0.0002%) and high (0.2%) inducer concentrations are combined. (**c**) Observed versus expected GFP expression values of all 300 pairwise (left) and 1000 triplet (right) genotypes. Points above the identity line have positive epistasis, whereas points below display negative epistasis. Data points represent the mean value of three biological replicates. Error bars for observed values (vertical) represent the standard deviation of three biological replicates. Error bars for expected values (horizontal) represent the calculated propagated errors from the errors of the single mutant genotype measurements (see Methods). (**d**) Different types of epistasis for all significant (2202 of 3900) pairwise (476 of 900) and triplet (1726 of 3000) combinations at the three inducer concentrations. (Left) Cases with all single mutant genotypes having a lower expression than the WT. (Middle) Cases with all single mutant genotypes having a higher expression than the WT. (Right) Cases of single mutant genotypes with mixed lower and higher expression than the WT. Percentages are based only on significantly epistatic genotype combinations and combined for pairwise and triplet combinations. sign: sign epistasis, magtd.: magnitude epistasis, RSE: reciprocal sign epistasis.

First, we calculated the observed fold change in fluorescence relative to the WT (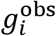 with *i* = 1, …, 30) for each of the 30 single mutant genotypes at each inducer concentration (**Extended figure 2.1a**). Under the multiplicative (log-additive) model, for a pairwise combination of mutations (*i*, *j*) we expect a fold change in fluorescence with respect to the mutant 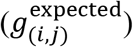 that follows,

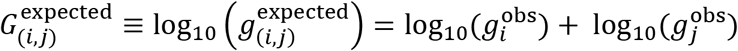

(**Extended figure 2.1b)**.

Similarly, for triplet mutants we have

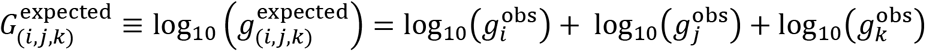

(**Extended figure 2.1c)**.

Next, we compared these expected values predicted from the single mutant genotypes with the actual observed GFP expression for pairwise or triplet combinations to calculate the magnitude of epistasis as

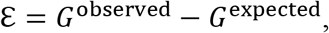

where *G*^observed^ is the logarithm of the observed fluorescence fold change of the mutant with respect to the wildtype 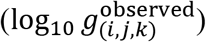. The magnitude and sign of the parameter ℇ measure the strength and sign of epistasis (**Figure 2a**). In particular, we defined epistasis values as significant if ℇ deviates from 0 with a false discovery rate (FDR) adjusted *p*-value of <0.05 (Domingo et al., 2018; Gao et al., 2010) **(Figure 2b**). From all measurements of pairwise combinations at all inducer concentrations, we found that 53% (476 of 900) resulted in significant epistasis (FDR adjusted *p*-value of <0.05) **(Figure 2c**). Similarly, 57% (1726 of 3000) of triplet combinations resulted in significant epistasis (FDR adjusted *p*-value of <0.05) (**Figure 2c**). Of the significant epistatic pairwise combinations, 82% were negative and only 18% positive, whereas for the triplet combination 88% were negative and only 12% positive (**Figure 2c**). Thus, most mutant genotype combinations resulted in lower GFP expression than expected.

Epistatic interactions can be classified depending on the phenotypic effects of each single mutant genotype and their combinations. For example, in magnitude epistasis the expression level associated to a genotype, but not its sign, changes with the genetic background more or less than would be expected under additivity. In contrast, if a genotype has the opposite effect when in combination with another genotype, i.e., it changes the sign of its relative effect, it is called sign epistasis. Reciprocal sign epistasis is a special case of sign epistasis, in which each single genotype has the opposite effect when combined with other genotypes (Domingo et al., 2019). Notably, most of the 2202 cases of significant epistasis for pairwise and triplet genotypes (n = 1230) could be attributed to negative reciprocal sign epistasis (RSE) (**Figure 2f**). In our case, single mutant genotypes had a higher gene expression than the WT network, but in combination they had a lower gene expression than any of the single mutant genotypes and the WT, resulting in negative RSE. These observations are in line with prior work in other systems, which uncovered epistasis amongst components that interact functionally (Lagator et al., 2016; Lagator et al., 2017a; Lagator et al., 2017b; Nghe et al., 2018) and physically (Anderson et al., 2015; Diss and Lehner, 2018). In sum, most interactions are epistatic in our system, with a preponderance of negative epistasis and negative reciprocal sign epistasis.

### Epistasis depends on the genetic background

Next, we compared the effects of the mutated nodes across the complete set of genetic backgrounds. For this we plotted the values of ℇ of all pairwise and triplet combinations for each of the 30 mutant genotypes separately and calculated the mean and variability of epistasis (**Figure 3**). Notably, we found that every genotype displayed both negative and positive epistasis, depending on the genetic background. Most genotypes have a negative mean value of ℇ, the exceptions being combinations of regulator-10, output-2 and output-10, whose mean ℇ was slightly positive (**Figure 3a and b**). To quantify the variability of epistasis we calculated the standard deviation of epistasis for each genotype (**Figure 3c**). The variability of epistasis decreased from sensor to regulator to output node, particularly at low and medium inducer concentrations, suggesting that the variability of ℇ was dependent on which network component was mutated (**Extended Figure 3.1**). In sum, these results show that epistasis is highly idiosyncratic in our system (Lyons et al., 2020) because the same mutation tends to have different effects in different genetic backgrounds.

**Figure 3.**
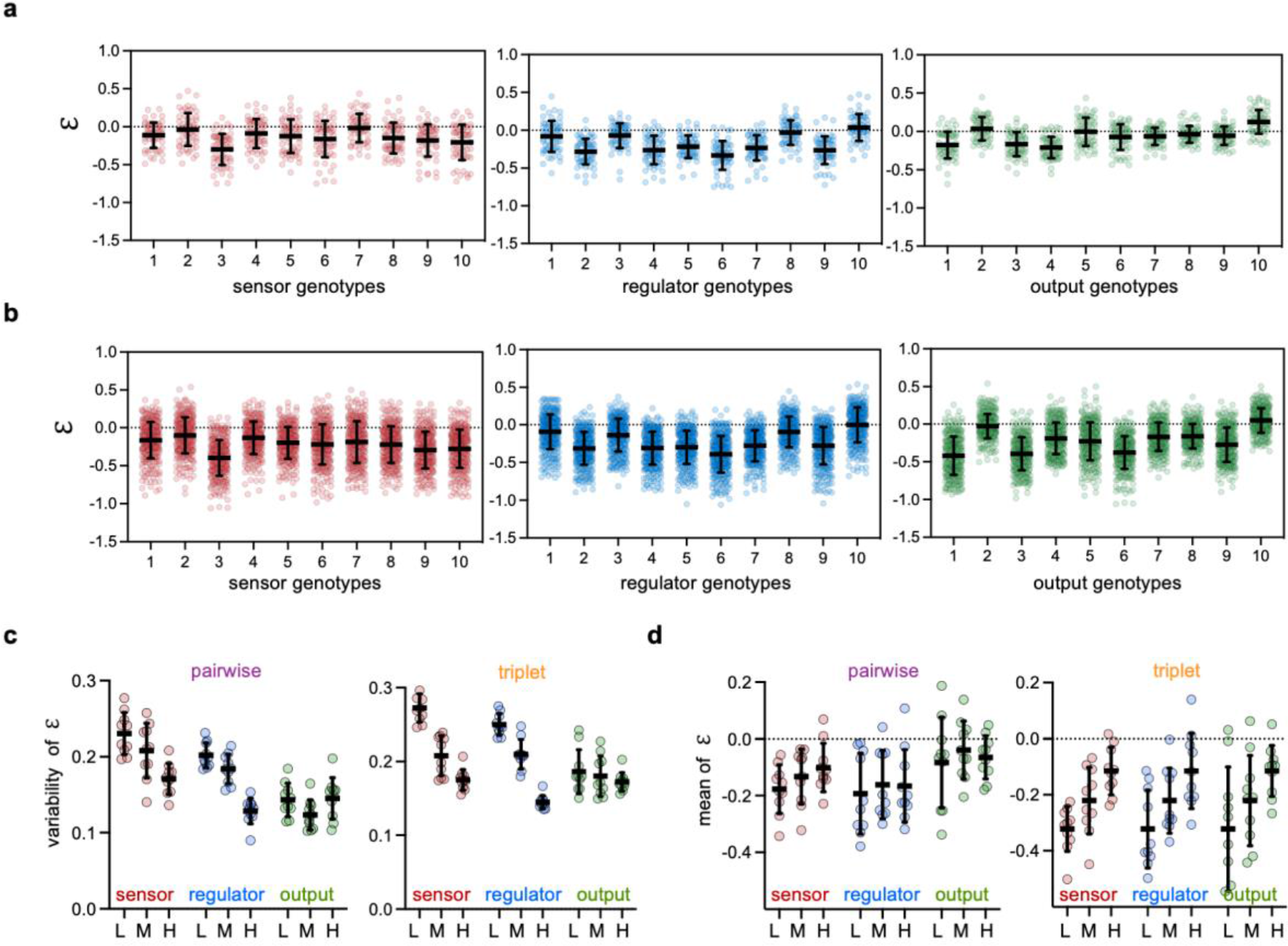
Sign and variability of epistasis varies with genetic background and inducer concentration. (**a**) Epistasis values for pairwise combinations of the 30 different mutant genotypes. Each genotype has 20 pairwise combinations with epistasis values at low (0%), medium (0.0002%) and high (0.2%) inducer concentrations giving rise to 60 values per genotype. Mean values represent the average epistasis of all 60 values and their variability is shown as the standard deviation. (**b**) Epistasis values for triplet combinations. Each genotype has 100 triplet combinations with epistasis values at low (0%), medium (0.0002%) and high (0.2%) inducer concentrations giving rise to 300 values per genotype. Mean values represent the average epistasis of all 300 values and their variability is shown as the standard deviation. (**c**) Variability of epistasis defined as the standard deviation of epistasis for all ten genotypes per node for pairwise (left) and triplet (right) combinations at low L (0%), medium M (0.0002%) and high H (0.2%) inducer concentrations. Horizontal bars represent the mean of the variability, and the error bars show the standard deviation. Statistical analysis is shown in **Extended Figure 3.1**. (**d**) Mean values of epistasis for all ten genotypes per node for pairwise (left) and triplet (right) combinations at low “L” (0%), medium “M” (0.0002%) and high “H” (0.2%) inducer concentrations. Horizontal bars represent the mean and the error bars show the variability as the standard deviation. Statistical analysis is shown in **Extended Figure 3.2**.

### Epistasis is inducer-dependent

Next, we asked if inducer concentrations influence the sign and variability of epistasis values. For this, we plotted the mean of epistasis values at low (0%), medium (0.0002%) and high (0.2%) inducer concentrations separately (**Figure 3d**). We found that the mean epistasis values increased, i.e. became more positive, with the inducer concentration for all three network nodes and this trend was more pronounced for triplet combinations (**Extended Figure 3.2** and **Extended Figure 3.3**).

To define significant inducer-dependent epistasis we performed a t-test with FDR adjusted p-values values at low, medium and high inducer concentration for each genotype of all pairwise and triplet combinations. Specifically, we defined a significant inducer-dependence of epistasis if the FDR q-value is <0.1 between any of the three inducer concentrations (**Extended Figure 3.4**). At this significance cut-off, 37% of pairwise (111 of 300) and 45% of triplet (447 of 1000) combinations exhibited significant inducer-dependent epistasis.

To further explore the inducer-dependence of epistasis in our dataset we explored how ℇ changed from low, medium to high inducer concentrations for each genotype combination. To do so, we classified the changes in epistasis along the inducer concentration into four categories, depending on which inducer concentration the epistasis value was highest or lowest (**Figure 4**): (a) with epistasis being highest at medium inducer concentrations; (b) with epistasis increasing with increasing inducer concentrations; (c) with epistasis decreasing with increasing inducer concentrations; (d) with epistasis being lowest at medium inducer concentrations. We quantified the distribution of pairwise and triplet combinations into these four categories. We found that out of all 300 pairwise combinations 33% fell into category (a), 31% into category (b), 18% each into categories (c) and (d). For the 1000 triplet combinations this distribution changed to 21% falling into category (a), 59% into category (b), 5% into category (c) and 15% in category (d). Thus, our results show that epistasis changes drastically with inducer concentration.

**Figure 4.**
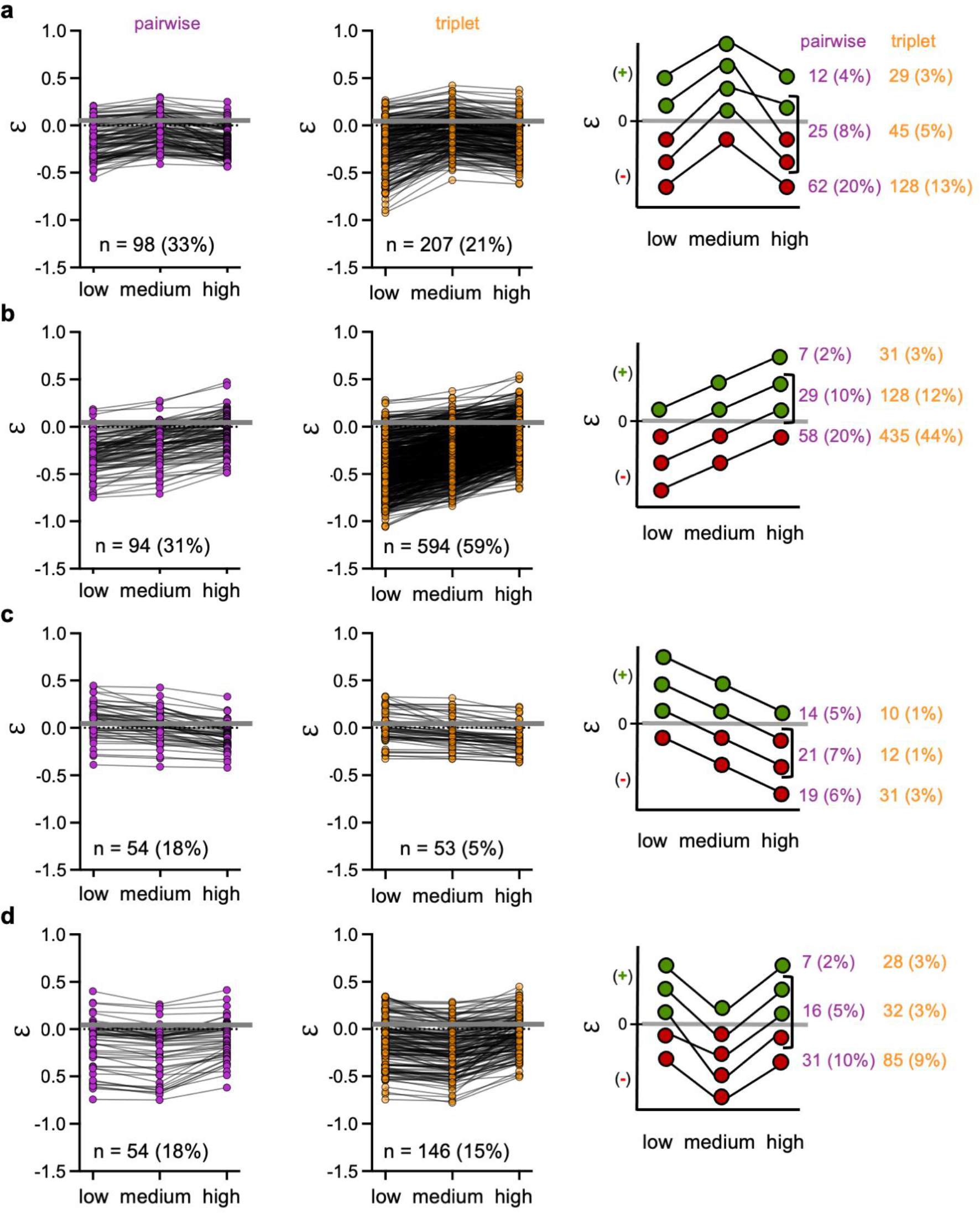
Epistasis changes with inducer concentrations. Epistasis along the inducer concentration gradient for pairwise combinations (**left**) and triplet combinations (**middle**). (**a**) Genotype combinations with epistasis highest at medium inducer concentration. (**b**) Genotype combinations with epistasis increasing with increasing inducer concentrations. (**c**) Genotype combinations with epistasis decreasing with increasing inducer concentrations. (**d**) Genotype combinations with epistasis lowest at medium inducer concentration. A quantification of the different scenarios of epistasis values changing or maintaining sign across inducer concentration is shown on the right.

Next, we asked if epistasis also switched sign between inducer concentrations. For this we sub-classified the four categories into three different scenarios: remaining negative, remaining positive or switching sign at different inducer concentrations (**Figure 4 – right part**). Overall, we found that 31% of the pairwise combinations switched sign along the inducer concentrations, and 13% remained always positive and 56% always negative. For the triplet combinations, we found that 21% switched sign and 10% remained always positive and 69% always negative. In sum, these results demonstrate that epistasis is environment-dependent in our system, changing in magnitude and sign along the inducer gradient.

### Inducer-dependent epistasis increases phenotypic diversity

We have shown that pairwise and higher-order epistasis depends on the inducer concentration in our system. Since the function of the network is to interpret an inducer gradient into a gene expression pattern, we next analyzed how this inducer-dependence of epistasis impacted the network’s pattern phenotype. To this end, we characterized the pattern phenotype of all single, pairwise and triplet genotypes. For this, we calculated the difference in gene expression between low and medium arabinose concentrations and between high and medium arabinose concentrations (**Figure 5a**) (Schaerli et al., 2018). This approach allows for a simple visualization on a cartesian plot and classification of pattern phenotypes into four categories for each one of the four quadrants: increase (Q1), anti-stripe (Q2), decrease (Q3) and stripe (Q4). Projections near the origin correspond to a constant expression phenotype.

**Figure 5.**
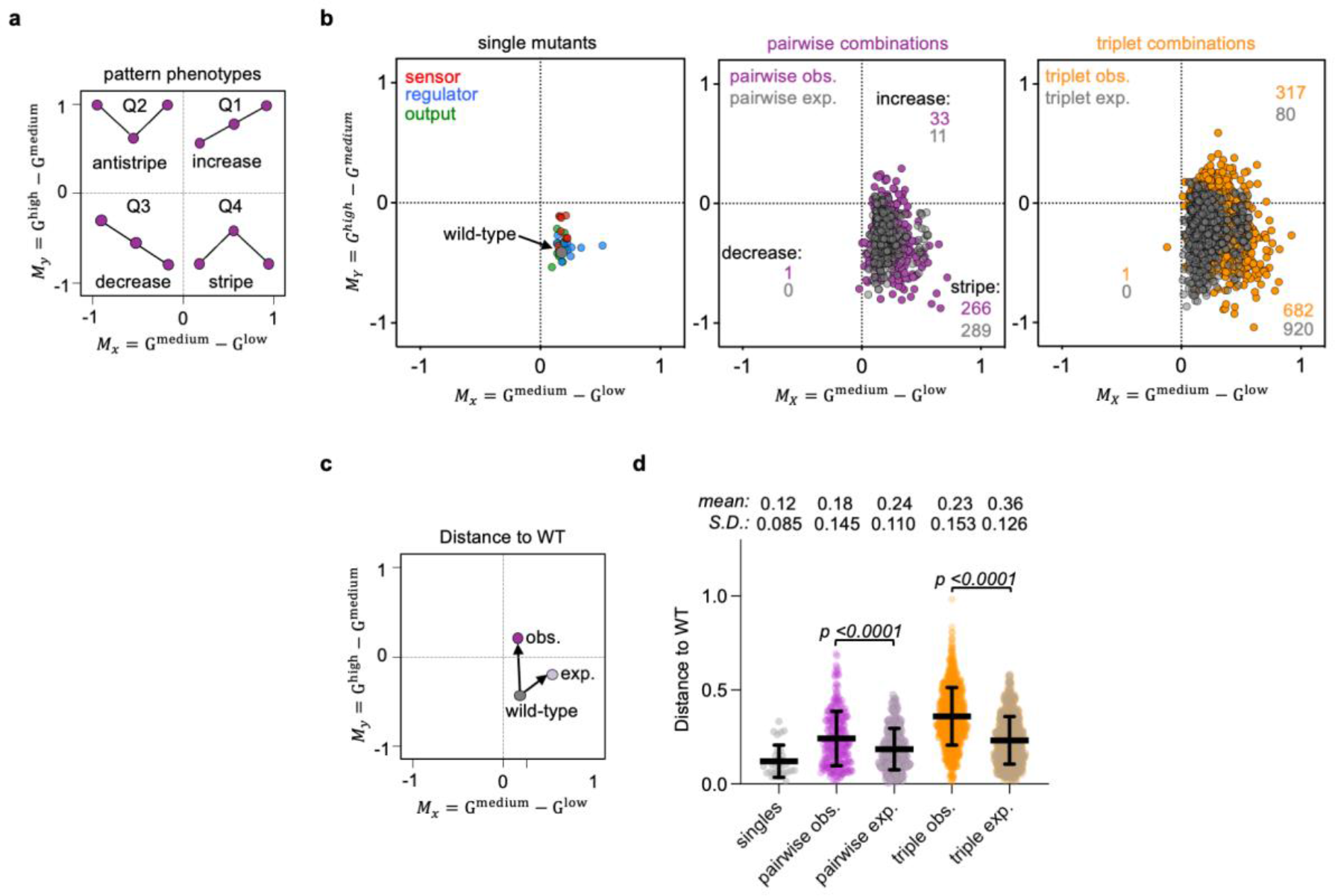
Pattern phenotypes. (**a**) Projection of pattern phenotypes to two-dimensional coordinates using ratios of GFP expression between medium-low (X axis) and high-medium (Y axis) inducer concentrations. (**b)** Pattern phenotypes of 30 single mutant genotypes (left) and expected (grey) and observed (colored) pairwise (middle) and triplet (right) genotypes. (**c**) Schematic of distance for pattern phenotypes relative to the wild type in the two-dimensional space depicted with arrows and calculated as Euclidean distance. (**d**) Measure of distance of pattern phenotypes relative to the wild type. Statistical significance (p-values) was calculated using a two-tailed paired t-test (Wilcoxon test).

We first plotted the pattern phenotypes of the WT and all 30 single mutant genotypes (**Figure 5b, left**). As expected, they all fell into the stripe category (Q4). We further examined the pattern phenotypes of all 300 pairwise (**Figure 5b, middle**) and 1000 triplet (**Figure 5b, right**) genotypes. Based on the multiplicative model, most of the pairwise (n=289 of 300) and triplet (n = 920 of 1000) genotypes were also expected to display a stripe pattern phenotype, with few exceptions (11 pairwise and 80 triplet) that were expected to adopt an increase pattern phenotype. These cases were the result of combinations of single mutant genotypes with high GFP expression at high inducer concentrations (e.g., sensor-3, -7, -10 and regulator-5, -6) or low GFP expression at low inducer concentrations (e.g., regulator-1 and output-10) (**Extended Figure 1.2**). However, the observed phenotypes were more frequently non-stripe patterns than expected. Specifically, for pairwise and triplet combinations we observed 22 (11 expected vs 33 observed) and 237 (80 expected vs 317 observed) more increase pattern phenotypes, respectively (**Figure 4b**). In addition, observed pairwise and triplet combinations also each showed one decrease phenotype.

In addition to the changes of distribution into the pattern phenotype categories, we also observed that the pattern phenotypes within one category were more diverse than expected from the multiplicative model. For example, several genotypes had a stronger stripe pattern phenotype than the WT, i.e., a bigger difference between lowest and highest GFP expression levels. To quantify and compare this spread of pattern phenotypes, we calculated the Euclidean distance between the pattern of the WT and the pattern of each genotype (schematic **Figure 5c**). We found that for most genotypes the observed mean distance to the WT was significantly higher than expected, both for pairwise (0.24 vs. 0.18, paired *t-test p*-value <0.001) and triplet combinations (0.36 vs 0.23, paired *t-test p*-value <0.001) (**Figure 5d**). Thus, we conclude that environment-dependent epistasis causes a greater diversity of pattern phenotypes than would be expected if the mutations did not interact epistatically or depend on the environment.

## Discussion

Mutations in cis-regulatory regions can alter the spatiotemporal gene expression patterns of gene regulatory networks. Such alterations are often deleterious, resulting in developmental abnormalities, disease, or death (Schaub et al., 2012; Ward and Kellis, 2012). However, they are occasionally advantageous, as evidenced by their common implication in evolutionary adaptations and innovations (Johnson, 2017; Prud’homme et al., 2007). Here, we used the toolkit of synthetic biology to systematically interrogate how combinations of mutations in the cis-regulatory regions of a three-gene regulatory network interact to influence the network’s spatial pattern phenotype. We uncovered pervasive epistasis, particularly negative epistasis and negative reciprocal sign epistasis. Moreover, we found epistasis to be more common among triplet combinations than among pairwise combinations, and that epistasis depended on the environment, because it varied across the inducer gradient. Finally, we showed that this inducer-dependent epistasis resulted in a greater diversity of pattern phenotypes than would be expected if the mutations did not interact epistatically and their phenotypic effects did not depend on the environment.

Prior work has shown that epistasis can either constrain or facilitate the evolution of phenotypic diversity. For example, in individual macromolecules such as DNA, RNA, and proteins, reciprocal sign epistasis has been shown to constrain the evolution of phenotypic diversity by forming maladaptive valleys in fitness landscapes, which can trap evolving populations on suboptimal adaptive peaks and preclude the generation of further adaptive phenotypic variation (Payne and Wagner, 2019; Poelwijk et al., 2011). However, when such macromolecules interact, epistasis amongst mutations in the interacting components can alleviate the constraints of the individual components, facilitating the evolution of phenotypic diversity (Anderson et al., 2015; Lagator et al., 2017b). Lagator and colleagues (2017b) provide a representative example. They studied the phenotypic effects of mutations in the canonical Lambda bacteriophage switch, a regulatory network consisting of RNA polymerase, a transcriptional repressor, and a cis-regulatory element to which both proteins bind. Their study showed that the phenotypic variation induced by combinations of mutations in the three interacting components was greater than that induced by mutations in the individual components. Moreover, the amount of phenotypic variation brought forth by combinations of mutations in the interacting components was different in the presence and absence of the transcriptional repressor, thus revealing environment-dependent epistasis in this system (Lagator et al., 2017b). Our study complements and extends this work by studying higher-order, environment-dependent epistasis in a larger regulatory network in which the mutations can interact functionally, but not physically (since mutations are only in cis-regulatory regions), to influence phenotypic diversity in a spatial pattern phenotype.

Our experiments also shed light on the robustness of gene expression pattern phenotypes to recombination, and how recombination generates novel phenotypes in gene regulatory networks. Prior work with computational models of gene regulatory networks compared the pattern phenotypes of recombinant offspring derived from parental networks that have the same phenotype to the pattern phenotypes of mutated offspring derived from these same parents (Martin and Wagner, 2009). Recombination was far less likely to cause a change in pattern phenotype than mutation. For example, more than 90% of recombinant offspring that differed from their parent by one regulatory interaction preserved the parental phenotype, as compared to only ~75% of mutated offspring that differed from their parent by one regulatory interaction. These differences in the robustness of pattern phenotypes to recombination and mutation only increased as the difference in the number regulatory interactions between parent and offspring increased. Our study provides experimental support for these findings, at least qualitatively. Specifically, in our previous work (Schaerli et al., 2018), we found that 67% of mutations to our regulatory network resulted in a non-stripe phenotype, whereas here we found that only 31.8% and 34% of pairwise and triplet genotype combinations resulted in a non-stripe phenotype, respectively. Relative to mutation, this is a 2-fold reduction. Further, we found that when recombination does generate a novel phenotype, this can be partly explained by environment-dependent epistasis. Without such epistasis, recombination is far more likely to preserve the parental pattern phenotype, as evidenced by comparisons with our null model. Our study thus supports the theoretical prediction of a low cost to recombination in creating novel phenotypes in gene regulatory networks (Wagner, 2011) and highlights environment-dependent epistasis as one cause of phenotypic novelty in recombinant offspring.

The robustness of a gene regulatory network’s pattern phenotype to mutation and recombination facilitates so-called “system drift” (True and Haag, 2001), in which an evolving population accumulates diversity in the regulatory and coding regions of the networks’ constituent components without causing a change in phenotype. System drift can facilitate the evolution of phenotypic novelties, because the resulting genetic diversity may serve as the basis for subsequent mutations or recombination to bring forth novel phenotypes, or it may be revealed as phenotypic variation upon environmental change (Payne and Wagner, 2019). Modeling work has long suggested that gene regulatory networks are susceptible to system drift, because many different mutationally-connected networks have the same expression phenotype (Ciliberti et al., 2007; Cotterell and Sharpe, 2010; Crombach et al., 2016; Jaeger, 2018), and recent empirical work has demonstrated a role for system drift in the evolution of biofilm formation in the fungus *Candida albicans* (Nocedal et al., 2017). Our work bridges these theoretical and empirical studies, using synthetic gene regulatory networks to experimentally interrogate the phenotypic effects of mutation and recombination, confirming the susceptibility of regulatory networks to system drift and the constructive role of the resulting genetic diversity in the evolution of novel phenotypes. A key finding of our study is that such novelties are at least partly explained by epistatic interactions amongst network components, which can change in magnitude and sign along an environmental gradient, thus altering gene expression levels across the spatial domain.

Our finding that environment-dependent epistasis can cause novel pattern phenotypes in recombinant offspring is germane to a growing body of literature on hybrid incompatibilities in gene regulatory networks (Johnson and Porter, 2000, 2007; Khatri and Goldstein, 2015, 2019; Palmer and Feldman, 2009; Porter and Johnson, 2002; Schiffman and Ralph, 2022; Tulchinsky et al., 2014a; Tulchinsky et al., 2014b). These modeling studies highlight several factors that influence the evolution of such hybrid incompatibilities, including population structure (Porter and Johnson, 2002), population genetic conditions (Khatri and Goldstein, 2015, 2019), mutational target size (Tulchinsky et al., 2014a; Tulchinsky et al., 2014b), network topology (Palmer and Feldman, 2009), and whether selection is directional or stabilizing (Porter and Johnson, 2002). Under stabilizing selection, where system drift can occur, these models suggest that hybrid incompatibilities are most likely to arise when many network variants that have the same pattern phenotype are mutationally connected with one another, forming so-called “genotype networks” (Wagner, 2008) or “neutral networks” (Schuster et al., 1994), and when at least some mutations in network components interact epistatically. Genotype networks facilitate system drift, because an evolving population can spread across the network while preserving pattern phenotype (Wagner, 2008), whereas epistasis can create maladaptive “holes” in these networks, into which recombinant offspring may fall (Gavrilets, 1997). Our work here and in previous studies (Santos-Moreno et al., 2022; Schaerli et al., 2018; Schaerli et al., 2014) provides experimental support that gene regulatory networks form genotype networks, and demonstrates that epistasis is not only prevalent amongst mutations in network components, but also depends on the environment. Thus, in line with prior modeling, our results suggest that system drift is likely to result in hybrid incompatibilities in recombinant regulatory networks. Whereas we observe that most recombinant offspring preserve the parental stripe phenotype, we emphasize that the genetic diversity in our parental population is limited. Recombination amongst a more diverse pool of parental networks may reveal a larger fraction of non-stripe phenotypes.

In sum, we used the toolkit of synthetic biology to perform a systematic analysis of the combined effects of mutations that are individually phenotypically neutral on the pattern phenotype of a gene regulatory network, uncovering pervasive pairwise and higher-order epistasis that changed in magnitude and sign along an inducer gradient, thus giving rise to novel spatial gene expression patterns that in natural gene regulatory networks may cause hybrid incompatibilities or embody evolutionary innovations. Such environment-dependent epistasis therefore strongly influences the evolution of gene regulatory networks, with implications for our understanding of speciation and evolutionary novelty.

## Materials and Methods

### Materials

Chemicals and media components, unless stated otherwise, were purchased from Sigma-Aldrich.

#### Media

For cloning and precultures we used Luria–Bertani medium (LB: 10 g Bacto-tryptone, 5 g yeast extract, 10 g NaCl per 1 l) supplemented with the appropriate antibiotics (100 μg/ml ampicillin for pET plasmids (output), 30 μg /ml kanamycin for pCOLA plasmids (sensor) or 50 μg /ml spectinomycin for pCDF plasmids (regulator)). For all plate reader assays of the synthetic networks “Stripe Medium” (SM) was prepared as follows: LB ingredients were dissolved in ultrapure water (Roth), sterile filtered (22 μm filter pores company) and supplemented with sterile 0.4% (w/v) glucose, 100 μg/ml ampicillin, 30 μg /ml kanamycin and 50 μg /ml spectinomycin and 5 μM isopropyl b-D-1-thiogalactopyranoside (IPTG).

#### The synthetic regulatory network and selected mutants

Our model system, a synthetic stripe-forming regulatory network based on the incoherent feed-forward loop type 2 architecture, was constructed and characterized previously (Schaerli et al., 2014). Each of the three nodes is encoded on a separate plasmid and the WT sequences are available at the NCBI GenBank (https://www.ncbi.nlm.nih.gov/genbank/). The GenBank accession codes are KM229377 (sensor, pCOLA backbone, kanamycin resistance marker), KM229382 (regulator, pCDF backbone, spectinomycin resistance marker) and KM229387 (output, pET backbone, ampicillin resistance marker). For a functional network to generate an inducer-dependent gene expression pattern, all three plasmids need to be transformed into *E. coli* MKO1 cells (Kogenaru and Tans, 2014). Mutated networks were selected from a previous study, in which we introduced mutations in the promoter and operator region of each node (Schaerli et al., 2018). Each single mutant genotype contains only mutations in one of the three plasmids, whereas the other two plasmids did not contain mutations. **Extended figure 1.1** shows the nucleotide changes in promoter and operator regions of the 30 selected mutant genotypes.

#### Generating pairwise and triplet genotype combinations

Plasmids of selected mutants were extracted and purified using the QIAprep Spin Miniprep Kit (Qiagen) according to the manufactures protocol. Each mutant selected from our previous study (Schaerli et al., 2018) contains one mutated and two wild-type plasmids. To extract only the mutated plasmids, we first removed the wild-type plasmids with restriction digest, retransformed the single plasmids into *E. coli* NEB5α cells, plated and cultured the cells with the appropriate antibiotics, and extracted the single plasmids. XhoI was used to isolate pCOLA plasmids (sensor), AatII and XmaI was used to isolate regulator pCDF plasmids (regulator) and XbaI was used to isolate pET plasmids (output).

We transformed the selected ten sensor, ten regulator and ten output mutants in all possible pairwise and triplet combinations into chemical competent MK01 *E. coli* cells. To generate pairwise combinations we transformed the two mutant plasmids into competent cells that already contained the third WT plasmid. For the triplet combinations, we transformed the sensor and regulator mutant plasmids (10 × 10 = 100) into competent cells that already carried one of the ten output mutant plasmids. Combinatorial libraries were cultured overnight in 96-deep well plates using selective LB medium and stored in 96-well plates at −80° C as glycerol stocks. Each 96-well plate contained 50 mutant genotype combinations and three wells with the wild-type variant and a well containing only media for data normalization purposes as described below.

#### Fluorescence measurement at 16 inducer concentrations

We measured the pattern phenotype over a gradient of 16 inducer concentrations for the wild-type, 30 single and 40 selected pairwise and triplet genotypes. We performed the measurements as follows: Starting from a glycerol stock we inoculated three 5 ml cultures (LB medium containing 0.4% (w/v) glucose and antibiotics) for each genotype, which served as our biological replicates and were from this point on treated independently. They were cultured overnight at 37°C and 200 rpm shaking. The following morning, we inoculated a fresh culture of 5 mL with 200 μl of the overnight culture followed by incubation for 3 h at 37°C and 200 rpm shaking. From these pre-cultures we used 5 μl to inoculate 384-well plates (Sigma-Aldrich, Nunc®, flat-bottom) containing 55 ul of SM media per well with inducer concentrations of 0.2%, 0.1%, 0.05%, 0.025%, 0.0125%, 0.00625, 0.00313%, 0.0016%, 0.0008%, 0.0004%, 0.0002%, 0.0001%, 0.00005%, 0.000025%, 0.000012% (w/v) arabinose. Immediately after inoculation 384-well plates were covered with clear lids to reduce evaporation and loaded into plate readers (Biotek Synergy H1) and measured as described below.

#### Fluorescence measurement of libraries

Measurement of the complete pairwise and triplet mutant libraries were performed as follows: starting from a glycerol stock plate, we inoculated three 96-well plates, which served as our biological replicates and were from this point on treated independently. Each 96-well plate contained one well with media only and three wells with the wild-type network, which were used for data normalization as described below. Each well contained 120 μl stripe LB medium containing 0.4% (w/v) glucose and antibiotics and plates were cultured overnight at 37°C and 700 rpm (THERMOstar, BMG Labtech). From each overnight culture 5 μl were used to inoculate a 120 μl selective LB medium pre-culture, which was incubated for 3 h at 37°C and 700 rpm (THERMOstar, BMG Labtech). From the preculture plate we used 5 μl to inoculate each of four different wells of a 384-well plate (Sigma-Aldrich, Nunc®, flat-bottom) each containing 55 μl of SM containing:

1. 0% arabinose (“low”)
2. 0.0002% arabinose (“medium”)
3. 0.2% arabinose (“high”)
4. 0.2% arabinose + 700 μM IPTG (“metabolic load” control)

Position A1 of a 96-well plate was used to inoculate position A1 (low), A2 (medium), B1 (high) and B2 (metabolic load) of a 384 well plate. The pipetting steps to inoculate 384-well plates were carried out using a semi-automatic pipetting robot (Rainin Smart 96, Mettler Toledo). Immediately after inoculation 384-well plates were covered with clear lids to reduce evaporation and loaded into plate readers (Biotek Synergy H1) and measured as described below.

#### Plate reader assay and data normalization

Microplates (96-well and 384-well) were incubated in plate readers (Biotek Synergy H1) with clear lids to reduce evaporation and shaken continuously in double orbital mode at a 2 mm radius and monitored for cell growth (optical density at 600 nm) and green fluorescence (excitation: 485 nm, emission: 520 nm) every 10 minutes at 37°C. Approximately after 3 h GFP expression peaked and *E. coli* cells started to reach stationary phase. As described previously (**Schaerli 2014**), the time-point when the fluorescence of the WT network at the medium arabinose concentration (0.0002%) peaked was chosen for further analysis of all fluorescence measurements. Fluorescence measurements were corrected for media background fluorescence and variation in cell number by dividing fluorescence with absorbance values. To adjust for plate-to-plate variation, we first normalized the three replicate plates based on the average of the three wild-type replicates on each plate. We then calculated the mean and standard deviation from the replicates and normalized again using the wild-type values from each plate to adjust for variation between all plates. The final data represent the average of three replicates, independent cultures started from the same glycerol stock, and errors correspond to the standard deviation between replicates.

Replicate measurements were excluded if values of cell growth differed by >0.2 from the absorbance of the WT controls on the same plate for any of the four conditions or suffered from metabolic load (as described previously (Schaerli et al., 2018)). Measurements were repeated if more than two replicates failed. Twenty-one out of the 1300 genotypes miss one replicate measurement due to growth differences. Note that we did not observe any metabolic load for any of the genotypes and combinations.

Correlations of R^2^ between replicates and between measurements of three and 16 inducer concentrations were calculated with Prism (Version 9.4.0, GraphPad Software, LLC.) using linear regression.

#### Calculating and defining significant epistasis

Based on a multiplicative model of epistasis, we calculated epistasis for all pairwise and triplet combinations at three inducer concentrations (low (0%), medium (0.0002%) and high (0.2%)) (Lagator et al., 2017a; Lagator et al., 2017b; Li et al., 2019; New and Lehner, 2019). First, based on the normalized fluorescence values we calculated the relative fold change in fluorescence (*g*_*i*_) for each of the 30 single mutant genotypes at each inducer concentration with respect to wildtype,

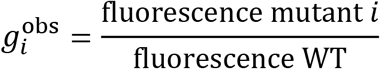

shown in **Extended figure 2.1a**. Under the multiplicative model, the expected relative changes of fluorescence are multiplicative. This allowed us to calculate the expected relative change in fluorescence of pairwise 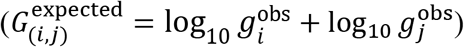 and triplet combinations 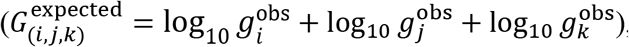, shown in **Extended figure 2.1b** and **c**. We next calculated the relative change in fluorescence for all measured 300 pairwise and 1000 triplet combinations at each inducer concentration,

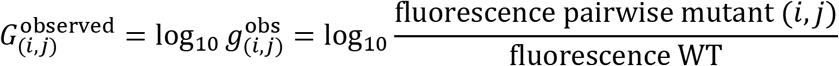

Similarly defined for the triplet combination, shown in **Extended figure 2.1b** and **c**. The definitions of *G* were used to calculate the epistasis for all pairwise or triplet genotype combinations,

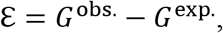

To define significant epistasis, we first calculated the uncertainty for each *G*^expected^ by propagating the log-transformed standard deviation (σ) from the single mutant measurements,

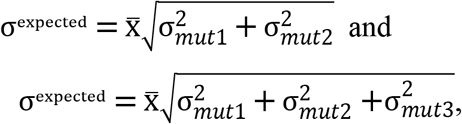

for pairwise and triplet combinations, respectively, with 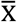 as the mean value of *G*^expected^. Using the mean of triplicate measurements and their SD or propagated errors, we then defined significant ℇ through series of t-tests (R function “tsum.test” with n.y and n.x = 3, alternative = “two.sided", var.equal = TRUE). The resulting *p*-values were then corrected for multiple testing using the “qvalue” package in R with its base parameters (reference). We defined epistasis values as significant if ℇ deviates from 0 with a false discovery rate (FDR) adjusted *p*-value of <0.05 (Domingo et al., 2018; Gao et al., 2010) **(Figure 2b**).

To define significant epistasis between low, medium inducer levels for each mutant genotype combination, we first propagated the error of observed and expected values as follows,

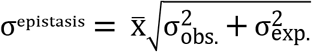

with 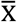 as the epistasis value and σ the standard deviation of *G*^observed^ and *G*^expected^, for each inducer concentration separately. Significant differences of epistasis between low, medium and high inducer levels were calculated with a series of t-tests (Welch’s test) and corrected for multiply testing with the Benjamini-Krieger-Yekutieli method in Prism (Version 9.4.0, GraphPad Software, LLC.). We defined significance values below a false discovery rate (FDR) q-value of <0.1.

#### Classifying types of epistasis

We classified epistatic interactions depending on the phenotypic effects of each single mutant genotype and their combinations into magnitude, sign and reciprocal sign epistasis (RSE) (Kemble et al., 2020). Their definition is schematically illustrated in **Figure 2d**. Briefly, magnitude epistasis is defined when the combined phenotypic effect of a genotype combination deviates from the expected effect but does not change the sign of the phenotypic effect of each single mutant. For instance, if two single mutant genotypes have a higher gene expression than the WT and their combination results in an even higher than expected gene expression, it is defined as positive magnitude epistasis. If their combined effect is lower than expected but remains higher than any of the two single mutant genotypes, this is defined as negative magnitude epistasis. A combination is classified as sign epistasis when the combined effect is lower than one of the two mutants, and thus changes the sign. For example, if two single mutant genotypes have a higher gene expression than the WT and their combination results in a value which is lower than one, but still higher than the other single mutant genotype. A special case of sign epistasis, reciprocal sign epistasis, occurs when the combined phenotypic effect has the opposite effect compared to any of the single mutant genotype’s effect. For example, RSE occurs when two single mutant genotypes have a higher gene expression than the WT, but their combined effect is lower than any single mutant genotype or even lower than the WT.

#### Pattern phenotype analysis

We visualized the pattern phenotypes on a cartesian plot (Schaerli et al., 2018). To this end, we calculated the difference in gene expression between low and medium arabinose concentrations for

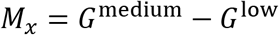

and between high and medium arabinose concentrations

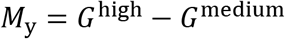

for all single mutant genotypes and observed and expected pairwise and triplet genotype combinations. The values of *M*_x_ and *M*_y_ for each genotype were then plotted as cartesian coordinates and classified into stripe (Q4), decrease (Q3), anti-stripe (Q2) and increase (Q1) pattern phenotypes, corresponding to the four quadrants. Projections near the origin correspond to a constant expression phenotype, i.e. a flat pattern phenotype.

To quantify and compare the spread of pattern phenotypes between observed and expected mutant genotypes, we calculated the Euclidean distance ‖*M*^exp^ − *M*^obs^‖ between the pattern of the WT and the pattern of each genotype as follows,

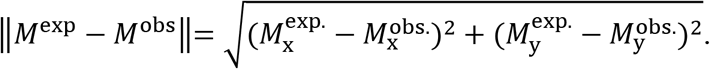

Significant differences between observed and expected *Euclidean Distance* values were calculated using a two-tailed paired t-test assuming no Gaussian distribution (Wilcoxon matched-pairs signed rank test) in Prism (Version 9.4.0, GraphPad Software, LLC.).

## Data availability

Source data of experiments are provided in **Source_Data**.

## Acknowledgments

RPC acknowledges the Life Sciences Department at Imperial College London. JLP acknowledges funding from the Swiss National Science Foundation (grants PP00P3_202672 and 310030_192541). YS also acknowledges funding from the Swiss National Science Foundation (grant 31003A_175608). We thank Florent Mazel, Dave W Anderson and Karol Buda for advice on statistical analysis.

## Competing interests

The authors declare that they have competing interests.

---

**Extended Figure 1.1.**
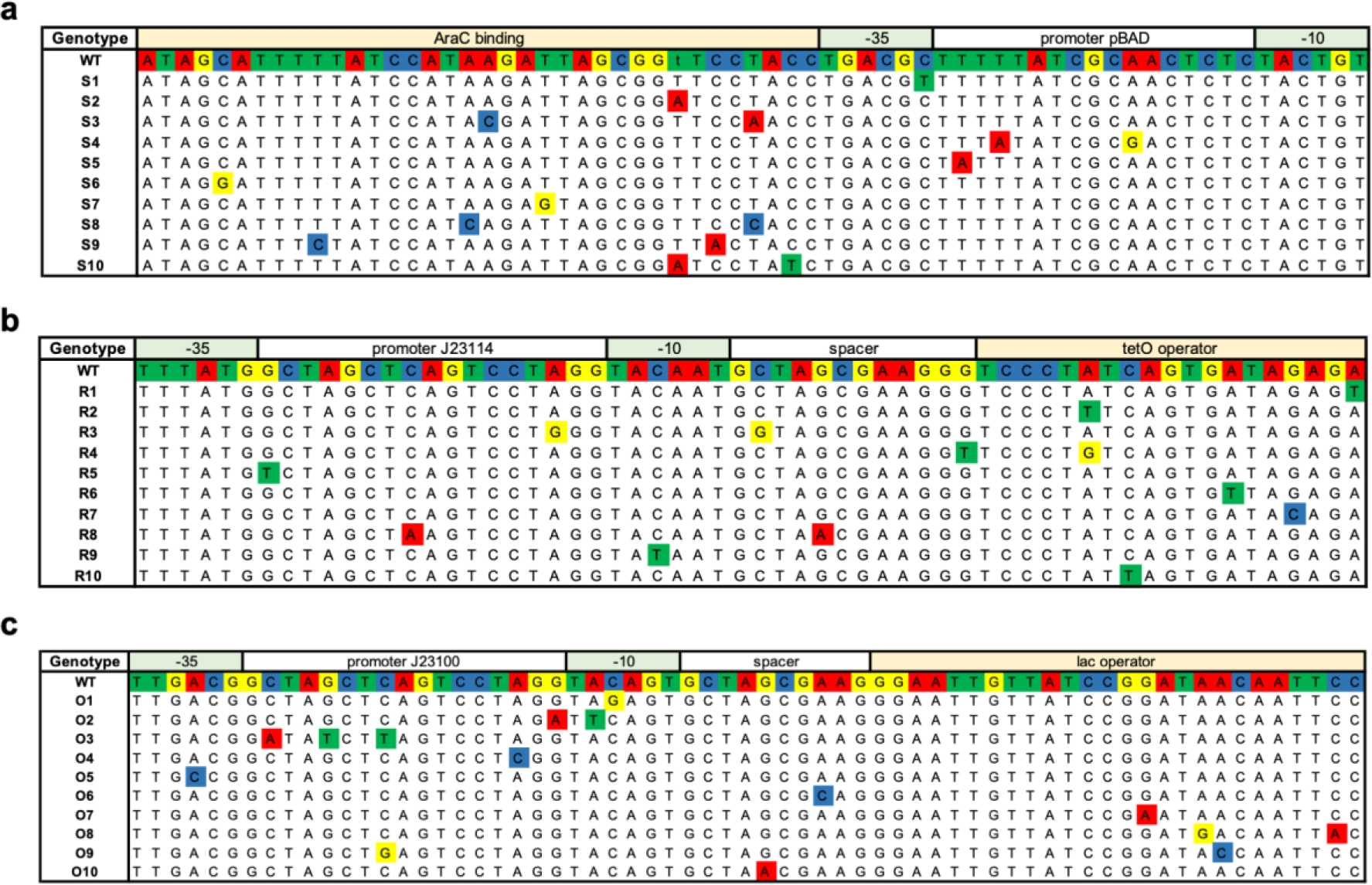
Nucleotide changes in promoter and operator regions of (**a**) sensor, (**b**) regulator and (**c**) output mutant genotypes.

**Extended Figure 1.2.**
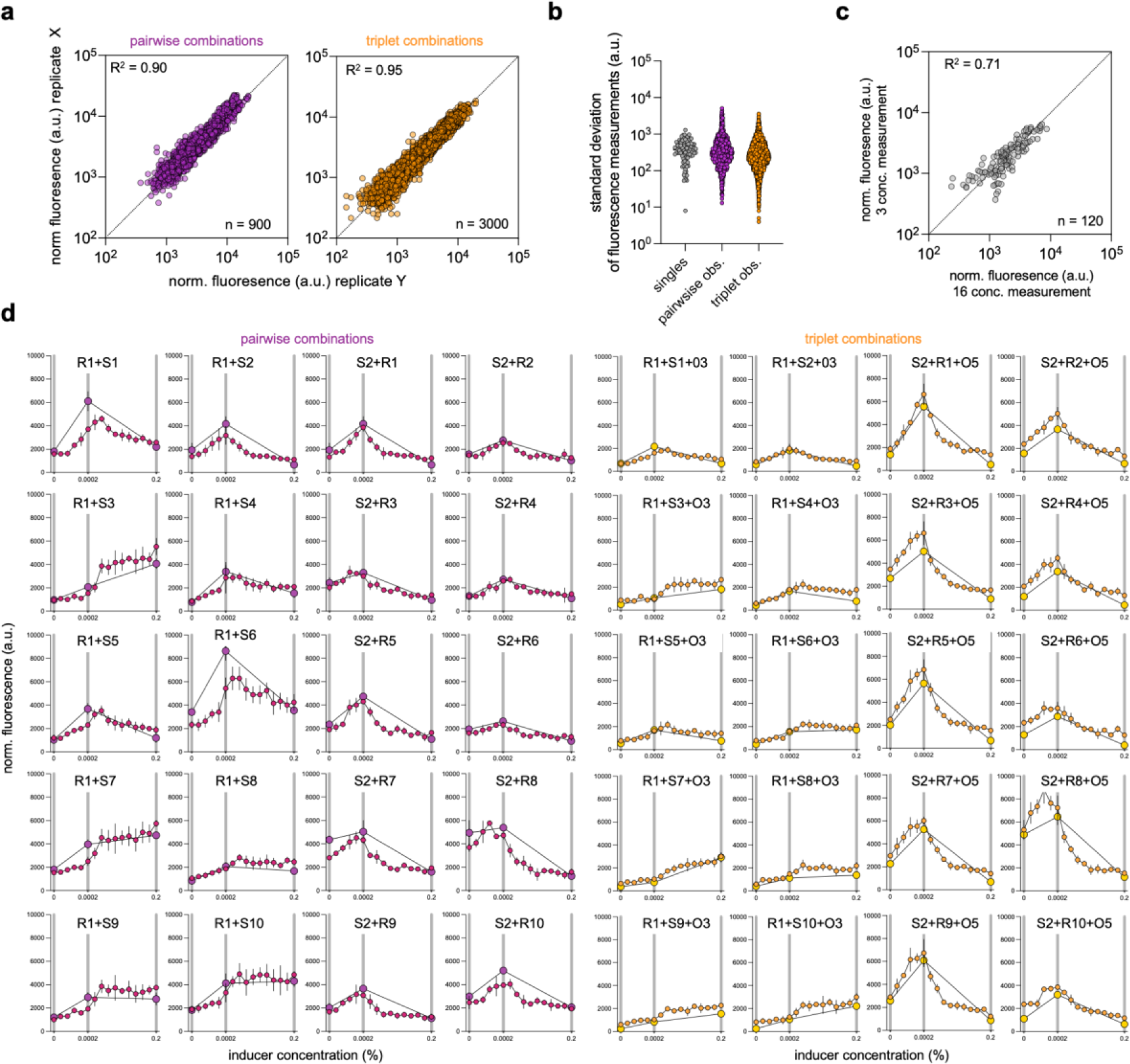
**(a)** Correlation between the three replicate measurements for all genotypes (left). X means replicate 1 or 2. Y means replicate 2 or 3. (**b**) Comparison of standard deviation values of fluorescence measurements between pairwise and triplet combinations. (**c**) Correlation between the two independent measurements of 40 selected genotypes shown in (d) at low, medium and high inducer concentrations. (**d**) Measured GFP expression pattern of the 40 selected pairwise (**left**) and triplet (**right**) genotypes. Each genotype was measured in triplicate at 3 (large points) and 16 (small points) inducer concentrations and the mean and standard deviation from three biological replicates are shown.

**Extended Figure 2.1.**
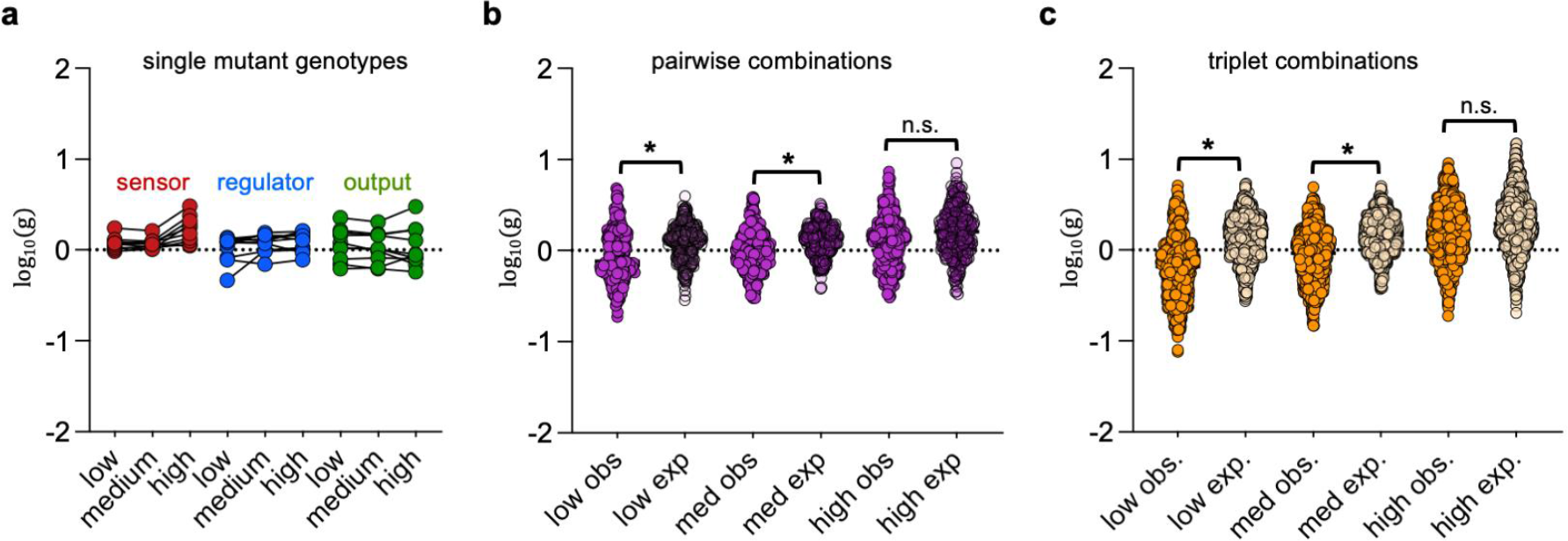
**(a)** Relative change in fluorescence (log_10_ (g)) of the 30 single mutant genotypes at low (0%), medium (0.0002%) and high (0.2%) inducer concentrations. Lines connect the same genotypes at different inducer concentrations. **(b)** Relative change in fluorescence (log_10_ (g)) of observed (obs.) and expected (exp.) pairwise combinations. **(c)** Relative change in fluorescence (log_10_ (g)) of observed (obs.) and expected (exp.) triplet combinations. Asterisks indicate significant difference of variability between the observed and expected relative fluorescence values (significance calculated using unpaired t-test (F test) with a of p-value <0.0001). n.s. means not significant with a p-value >0.0001.

**Extended Figure 3.1.**
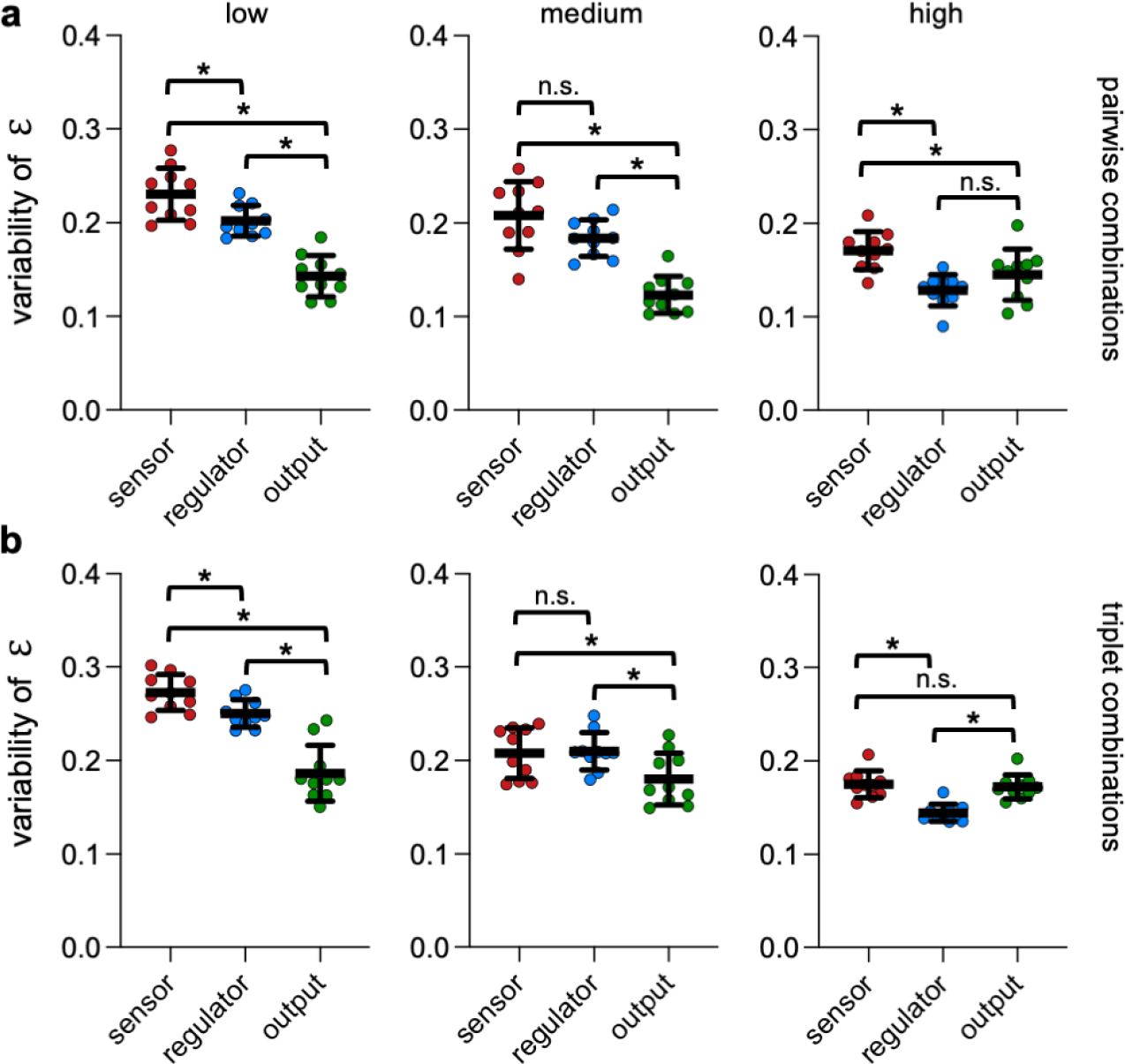
Statistical analysis for the variability of epistasis between regulatory nodes for (a) pairwise combinations and (b) triplet combinations at different inducer concentrations. Asterisk indicates significant difference of variability between the observed and expected relative fluorescence values (significance calculated using unpaired t-test with Welch’s correction. Asterisk indicates significant differences of the mean with a p-value <0.05). n.s. means not significant p-value >0.05.

**Extended Figure 3.2.**
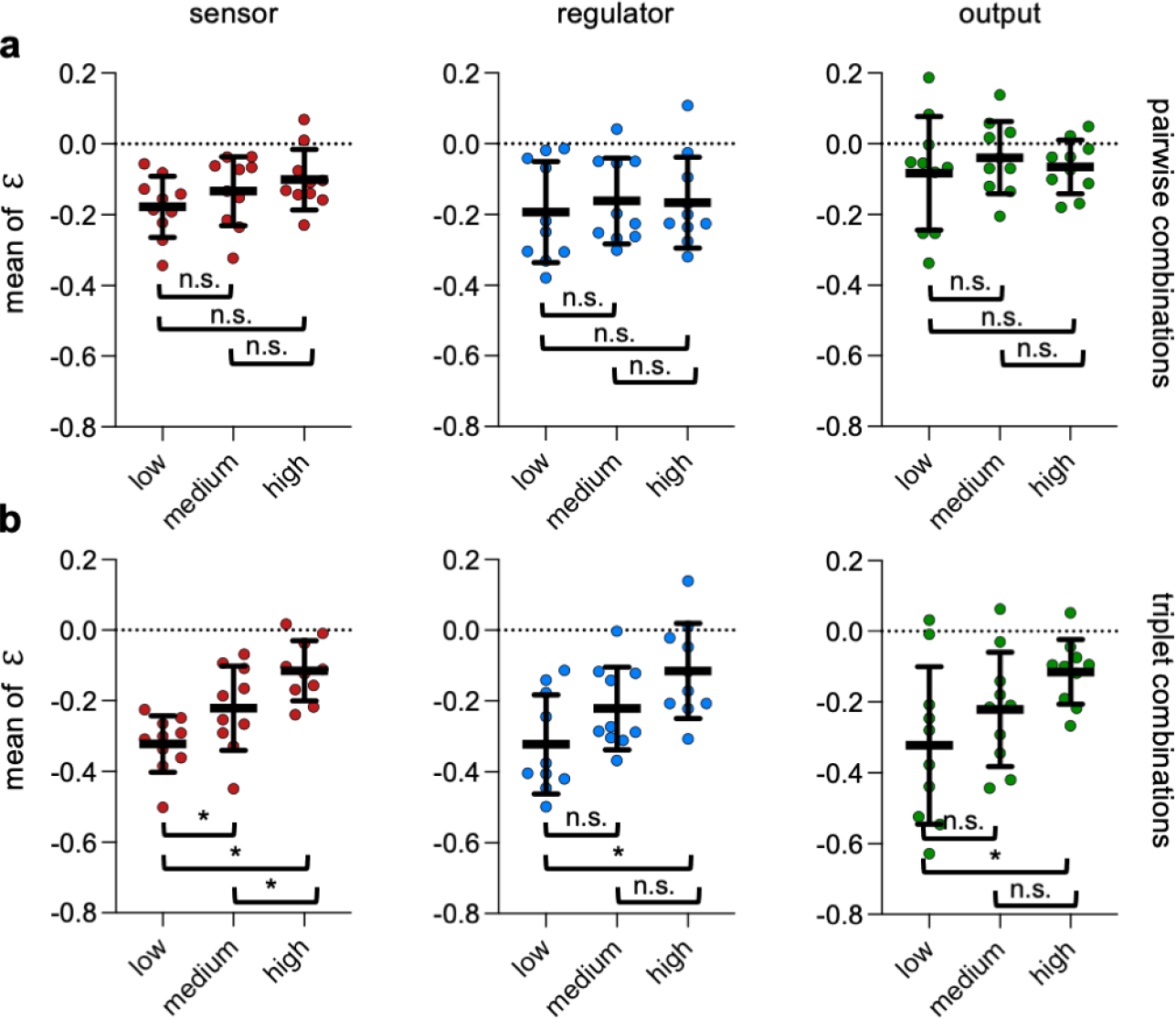
Statistical analysis for mean values of epistasis between different inducer concentrations for (a) pairwise combinations and (**b**) triplet combinations for the different regulatory nodes. Asterisk indicates significant difference of mean between the observed and expected relative fluorescence values (significance calculated using unpaired t-test with Welch’s correction). Asterisk indicates significant differences of the mean with p-values <0.05 and n.s. means not significant with p-values >0.05.

**Extended Figure 3.3.**
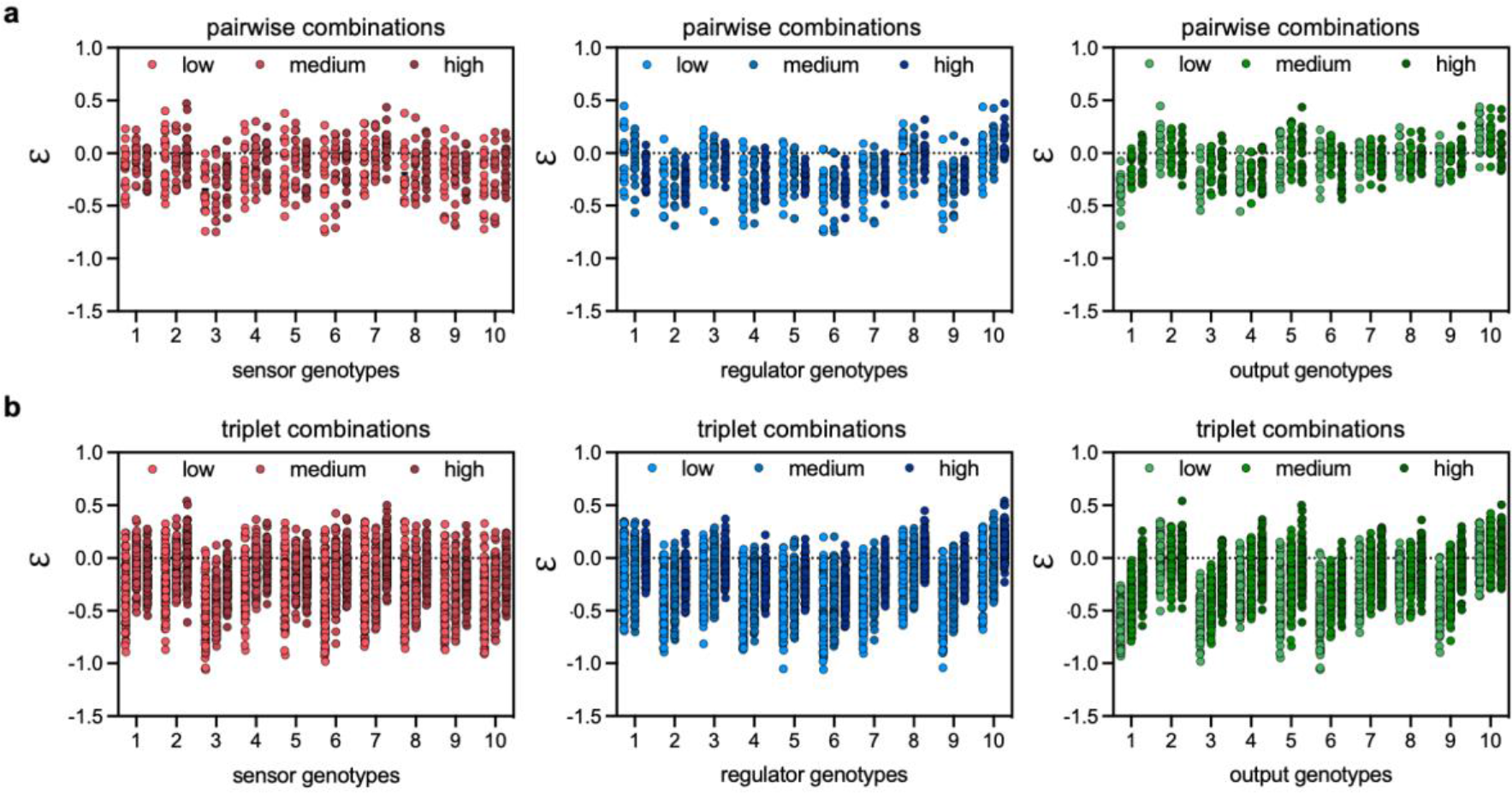
Epistasis is inducer-dependent. Epistasis values for each mutant genotype and inducer concentration for (**a**) pairwise combinations and (**b**) triplet combinations.

**Extended Figure 3.4.**
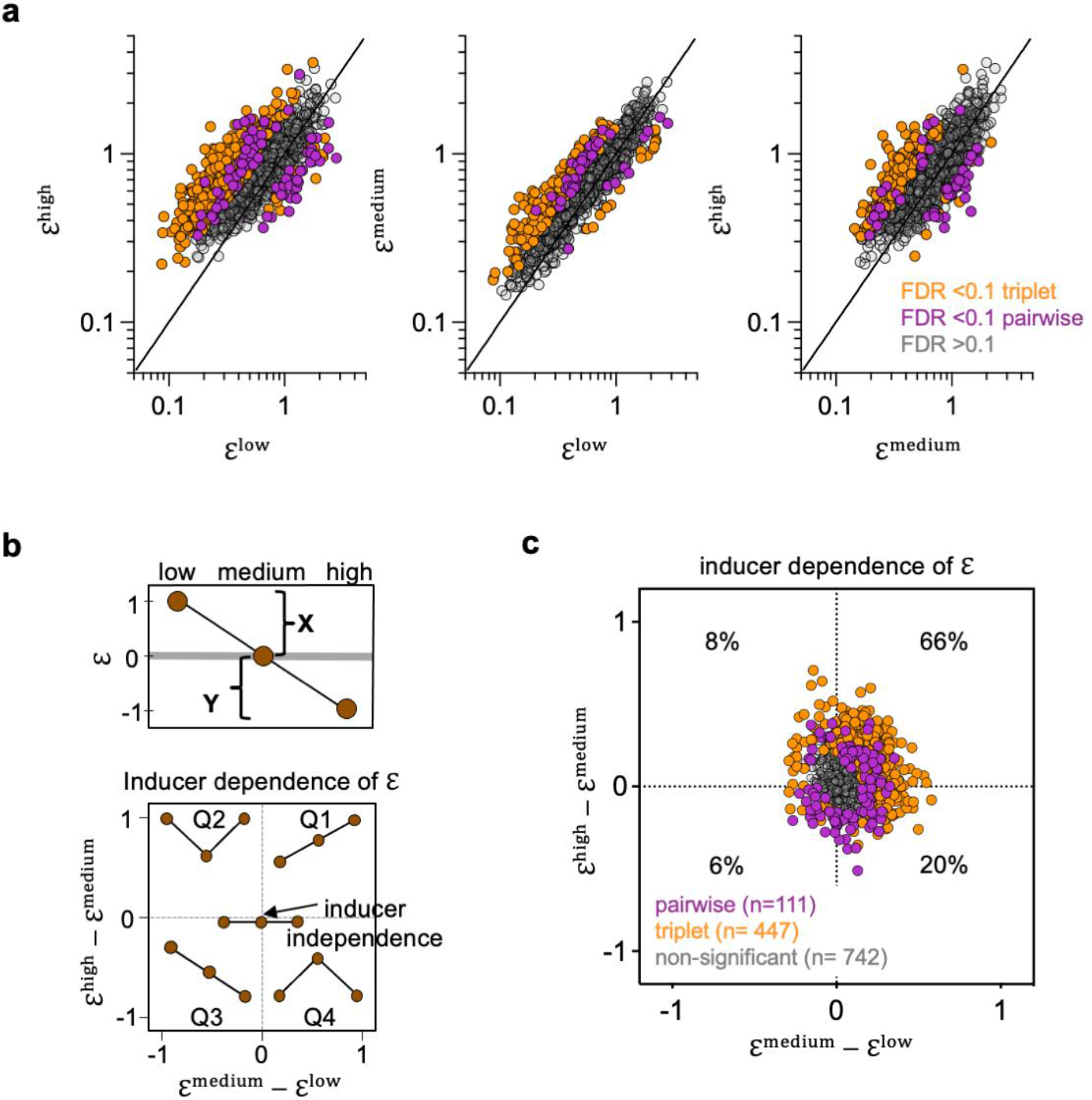
Defining significant inducer-dependent epistasis. (**a**) Defining significant inducer-dependent epistasis of all 300 pairwise (purple) and 1000 triplet (orange) genotype combinations. Significance was calculated with a series of t-tests (Welch’s test) with FDR correction (n = 3900, Benjamini-Krieger-Yekutieli method) with significant values (FDR q-value <0.1) in colour and non-significant values (FDR q-value >0.1) in grey. At this significance cut-off, 37% of pairwise (111 of 300) and 45% of triplet (447 of 1000) combinations exhibited significant inducer-dependent epistasis. (**b**) Projection of inducer-dependence of epistasis (ℇ) to two-dimensional coordinates using ratios of ℇ between medium-low (X axis) and high-medium (Y axis) inducer concentrations. (**b)** Genotypes with significant inducer-dependence are shown in color (pairwise in purple and triplet combinations in orange) and genotypes with non-significant inducer dependence are shown in grey.

## Notes

### Competing Interest Statement

The authors have declared no competing interest.

